# *ML-morph*: A Fast, Accurate and General Approach for Automated Detection and Landmarking of Biological Structures in Images

**DOI:** 10.1101/769075

**Authors:** Arthur Porto, Kjetil L. Voje

## Abstract

1. Morphometrics has become an indispensable component of the statistical analysis of size and shape variation in biological structures. Morphometric data has traditionally been gathered through low-throughput manual landmark annotation, which represents a significant bottleneck for morphometric-based phenomics. Here we propose a machine-learning-based high-throughput pipeline to collect high-dimensional morphometric data in images of semi rigid biological structures.
2. The proposed framework has four main strengths. First, it allows for dense phenotyping with minimal impact on specimens. Second, it presents landmarking accuracy comparable to manual annotators, when applied to standardized datasets. Third, it performs data collection at speeds several orders of magnitude higher than manual annotators. And finally, it is of general applicability (i.e., not tied to a specific study system).
3. State-of-the-art validation procedures show that the method achieves low error levels when applied to three morphometric datasets of increasing complexity, with error varying from 0.5% to 2% of the structure’s length in the automated placement of landmarks. As a benchmark for the speed of the entire automated landmarking pipeline, our framework places 23 landmarks on 13,686 objects (zooids) detected in 1684 pictures of fossil bryozoans in 3.12 minutes using a personal computer.
4. The proposed machine-learning-based phenotyping pipeline can greatly increase the scale, reproducibility and speed of data collection within biological research. To aid the use of the framework, we have developed a file conversion algorithm that can be used to leverage current morphometric datasets for automation, allowing the entire procedure, from model training all the way to prediction, to be performed in a matter of hours.

## 1 Introduction

In the past 20 years, genomics has revolutionized our understanding of biology, leading to the discovery of a wealth of novel phenomena (Koboldt et al, 2013). These discoveries were only possible through the development of a technological infrastructure that allowed us to acquire genomic information at a large scale (Schuster, 2007). While many researchers have argued that phenomics - large scale phenotyping - will bring about a similar revolution in biology, most approaches for collecting high-throughput phenotypic data developed so far are system-specific and difficult to generalize (see Kristensen et al., 2008; Falkingham, 2012; Boyer et al., 2011; Hsiang et al., 2018; Manacorda and Asurmendi, 2018), with the notable exception of sub-cellular phenotypes (e.g., Clish, 2015). As a consequence, for most study systems, phenotyping methods remain low-throughput and can only be applied on a small scale (Houle, Govindaraju, and Omholt, 2010).

These limitations have important consequences for biological research. First, the majority of the history of life remains inaccessible to genomic information, due to DNA degradation (Allentoft et al., 2012), and therefore needs to be integrated into current evolutionary theory through its phenotype. Second, even with the advances that came with genomics, we still cannot understand a wealth of important biological phenomena, such as disease and evolutionary fitness, suggesting that we have not been able to fully characterize such phenomena (Houle, Govindaraju, and Omholt, 2010). Third, phenomic and genomic research are largely synergistic, and the end product of being able to characterize both aspects of biological variation simultaneously is likely to increase the power of each approach (Houle, Govindaraju, and Omholt, 2010).

The low-throughput nature of current phenotyping methods is particularly problematic in morphometrics. Morphometrics has become an indispensable component of statistical analyses of size and shape variation in biological structures, with thousands of papers that make use of it being published every year (Hook et al., 2018). In the last two decades, with the growth of geometric morphometrics, geneticists, evolutionary biologists, ecologists and paleobiologists have accumulated dense landmark datasets, often collected in thousands of specimen images. Despite considerable progress in multivariate statistical analyses of morphometric data (Adams et al., 2016) and attempts of incorporating it into phylogenies (Parins-Fukuchi, 2018), we are still far from a comprehensive understanding of multivariate patterns of morphometric variation. Understanding multivariate patterns of variation requires large sample sizes (Grabowski and Porto, 2017). One of the main impediments to the acquisition of large landmark datasets is the manual collection of landmark data, which is both time and labor intensive. Given the recent explosion of semi-landmark use in morphometrics (Watanabe, 2018), data collection time has only increased. Depending on the number of landmarks and the necessary steps to prepare a specimen for landmark data collection, this manual annotation can take months, if not years.

A promising way to collect high-throughput phenotypic data is to automate landmark data collection using computer vision techniques (e.g., Manacorda and Asurmendi, 2018). Automated landmarking has become the gold standard in human facial landmarking for both biomedicine (Porto et al., 2019) and, more notoriously, social networking websites and software developed for mobile phones (Kazemi and Sullivan, 2014), but its application in geometric morphometrics has remained restricted (Manacorda and Asurmendi, 2018; Vandaele et al., 2018). However, the explosion in machine learning algorithms for computer vision represents an important technological leap, which lays the foundation for the development of general methods of high-throughput high-dimensional morphometrics (Kazemi and Sullivan, 2014; He et al., 2017; Voulodimos et al., 2018).

Here we develop a supervised learning-based phenotyping pipeline (ML-morph) to collect high-dimensional morphometric data in images of semi-rigid biological structures. This pipeline is based on adapting methods currently being used in computer vision research to morphometrics and allows for dense and accurate landmarking at low cost, high speed, and with minimal impact on specimens. Since morphometrics has traditionally relied on specialized software (Rohlf, 2006) and R packages (Adams et al., 2016), while most computer vision libraries are implemented in Python or C++(e.g., King, 2009), we also develop file conversion algorithms to leverage current morphometric datasets for training the machine-learning phenotyping pipeline. Development of a machine-learning infrastructure will enable biologists to get the necessary data to investigate and tackle important theoretical and empirical challenges within the field, and will greatly increase the scale, reproducibility and reach of biological research.

## 2 Methods

### 2.1 Description of the framework

We approach the problem of automated landmark detection using a supervised learning approach. In this approach, automated landmarking is performed by a combination of object detection (Dalal and Triggs, 2005) followed by shape prediction (Kazemi and Sullivan, 2014). In other words, our models are trained to (i) predict the location of a biological structure we intend to landmark in images (object detection) and, to (ii) predict the shape of each detected structure (i.e. annotate landmarks). Object detection and shape prediction models are trained using a dataset of manually annotated images (see Image annotation). In the following sections, we present and describe in detail the training process that can be used to generate detectors and predictors, the set of images and landmarks used to explore the performance of the method, and finally the metrics employed to validate the approach and to quantify how reliable it is in comparison with system-specific tools that have been published in the literature.

### 2.2 Sample

We tested the below proposed framework on three morphometric datasets (fly wings, sea basses and bryozoan colonies) with different levels of complexity. Figure 1 shows the landmarks we analyzed in each of the three datasets.

**Figure 1:**
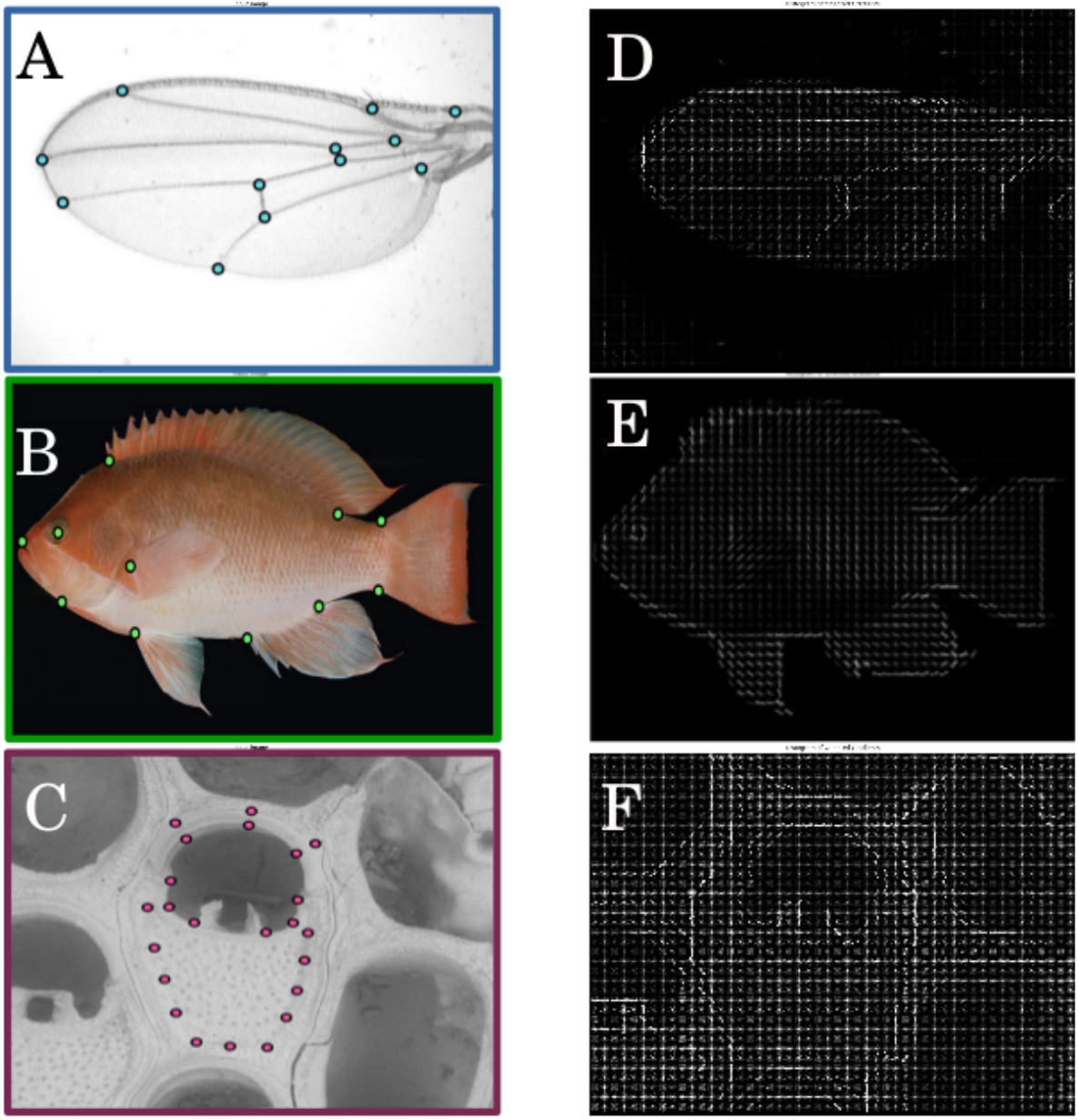
Landmark datasets and corresponding histogram of gradient (HOG) features. Three landmark datasets of increased difficulty were used to test the performance of the automated landmarking framework. Our framework uses HOG features (second column) to detect objects within an image (first column). (A) Drosophilid wing – (B) Sea bass (genus *Pseudanthias*) - (C) Bryozoan colony (*Steginoporella magnifica*) (D-F) Histogram of gradient features of images A-C.

#### Low complexity image set

The drosophilid wing is a structure commonly used in morphometric studies in ecology and evolution (e.g., Houle et al., 2003). We consider it to be a relatively simple dataset to develop automated landmarking algorithms for, in large part because: (1) the structure is translucent, creating clear contrast between structure and background, (2) the structure itself is highly conserved across species (Houle et al., 2017), and (3) landmarks are positioned in clear intersections of wing veins. Indeed, image analysis has already been automated for drosophilid wings using a B-spline model of wing shape (Houle et al., 2003), which makes this an ideal structure to investigate the performance of our machine-learning phenotyping pipeline. More specifically, we obtained a total of 280 images of drosophilid wings from Morphbank (Supporting Information 1) and placed a total of 12 landmarks in each image, following (Houle et al., 2003). Images have 632×480 pixels resolution and have been randomly selected from a larger pool of images. The reason to limit the number of images is to mimic the size of datasets that are usually used in morphometric studies. Figure 1A shows all 12 landmarks collected in each specimen.

#### Intermediate complexity image set

We analyzed 180 lateral images of sea basses belonging to genus *Pseudanthias* (Randall, 1997). This genus was chosen because it is composed by a large array of species with diverse morphologies, not only in terms of body shape but also in terms of coloration. These images have been captured by Dr John E. Randall and are deposited at the Bernice Pauahi Bishop Museum (Honolulu, Hawaii). They represent an intermediate level of complexity for the pipeline because the morphological diversity is much larger among these organisms than what is observed in drosophilids. Background color and image resolution are also more varied across image files compared to the drosophilids wings. Following the approach used for drosophilids, a total of 12 landmarks were manually annotated in each fish by the same expert (Figure 1B).

#### High complexity image set

Our high complexity dataset was collected in-house (UiO) and consists of 400 scanning electron microscope (TM-4000; Hitachi, Tokyo, Japan) images of bryozoan colonies belonging to the species *Steginoporella magnifica*. This data has been acquired as part of a fossil time-series project and is composed exclusively of Plio-Pleistocene fossil specimens collected from the Wanganui basin, New Zealand (Carter and Naish, 1998; Liow et al., 2017). We consider these pictures more complex than the fish and wing pictures, as the material contains colonies with different levels of fossil degradation. In addition, colonies contain multiple zooids with extreme plasticity in morphological traits, requiring the identification of individual zooids from whole-colony image data, while the fish and fly wing pictures contained only one specimen per image. We imaged all S. magnifica colonies at 1280×960 resolution and annotated individual zooids for a total of 23 landmarks (Figure 1C).

### 2.3 Image annotation – Defining ground truths

An essential component of supervised learning is image annotation. In our case, prior to model fitting, we annotated all images belonging to the three datasets using both bounding box annotations and individual landmark XY coordinates using *imglab* (King, 2009). Bounding box annotations are used to locate objects within images in order to train object detectors, while XY coordinates are used as landmark positional ground truths by the shape prediction part of the pipeline. A total of 10% of all images from each of the three datasets were annotated twice (3 months later), allowing us to estimate intra-observer measurement error. The intra-observer error is a key parameter, as it limits the maximum accuracy of any machine-learning algorithm. Here, we use intra-observer error as the theoretical minimum error our pipeline could hope to achieve.

Once annotated, we split all three datasets into training and validation sets, representing 80% and 20% of the images, respectively. We used the training set to fit the parameters of the object detectors and shape predictors. We then used the validation set to evaluate the performance of these algorithms. We employ metrics that are standard in problems of such kind to evaluate the performance of our models (see *Testing* for details).

### 2.4 Training object detectors

The first step in any computer vision algorithm for automated landmarking is the detection of the presence and position of the structure of the interest within the image. For example, if we want to identify landmarks in still images of drosophilid wings, we need the ability to identify the number and position of wings within an image. While a large number of object detection algorithms have been published in the past few years (King, 2009; Viola and Jones, 2001; Girshick, 2015; Ren et al., 2015 to name a notable few), the basic procedure for object detection remains largely the same. First, a set of positive (object, e.g., wing) and negative (not object, e.g, background) image windows are produced based on annotated training images and then a binary classifier is trained based on these windows. Finally, this classifier is tested on images in which the main model has not been trained on.

While the current state of the art in object detection relies on convolutional neural networks (CNN, e.g., Ren et al., 2015), these can be overpowered for standard biological applications. They also require large training samples (one to two orders of magnitude higher than the one proposed here), more fine-tuning of training parameters, specialized hardware (e.g., GPU), and usually perform at lower image processing rates (images per second) than simpler models (Suleiman et al., 2017).

Our implementation of object detection is based on histogram of oriented gradients (HOG) features within a sliding window framework (Dalal and Triggs, 2005; Figure 2). In brief, based on each training image, we randomly extract image regions that contain and do not contain the object of interest using bounding box annotations. From each randomly extracted region, we extract HOG features and create a training set. We then train a structural support vector machine (SVM) classifier to classify images according to these two labels (object vs. no object). In order to perform detection on the test set, we scan this classifier over an image pyramid using a sliding window, and, whenever a certain window passes a threshold test, it is output as being an object, after non-maximum suppression is performed. HOG feature generation follows the general approach developed by (Dalal and Triggs, 2005). We start by dividing each image into 8×8 cells and compute the gradient vector at each pixel. Following this step, we end up with a 64 (8×8) gradient vectors for each cell, which are then represented as histograms. These histograms compress the information in each cell by splitting that information into angular bins, where each bin corresponds to a gradient direction (20 degrees each). We then normalize the gradients using block normalization. During training, two parameters of SVMs require particular attention: the soft margin parameter C and the insensitivity zone (*ϵ*) (Kecman, 2001, page 182-183). The C parameter regulates the size of the margin, which is the distance between the hyperplane that separates the two classes and the closest data point. The *ϵ* parameter, on the other hand, regulates the penalty associated with errors in classification. In order to evaluate the impact various C and *ϵ* parameters (the training parameters) have on the performance of the final model, we performed an exhaustive grid search, in which a model is trained for each possible combination of hyperparameters, given a certain parameter range. In our case, we varied both the SVM C (from 1 to 7) and *ϵ* parameters (from 10^*−*2^ to 10^*−*4^). The combination of hyperparameters that results in the best model performance is then used in the final model, preventing errors that are incurred by overfitting.

**Figure 2:**
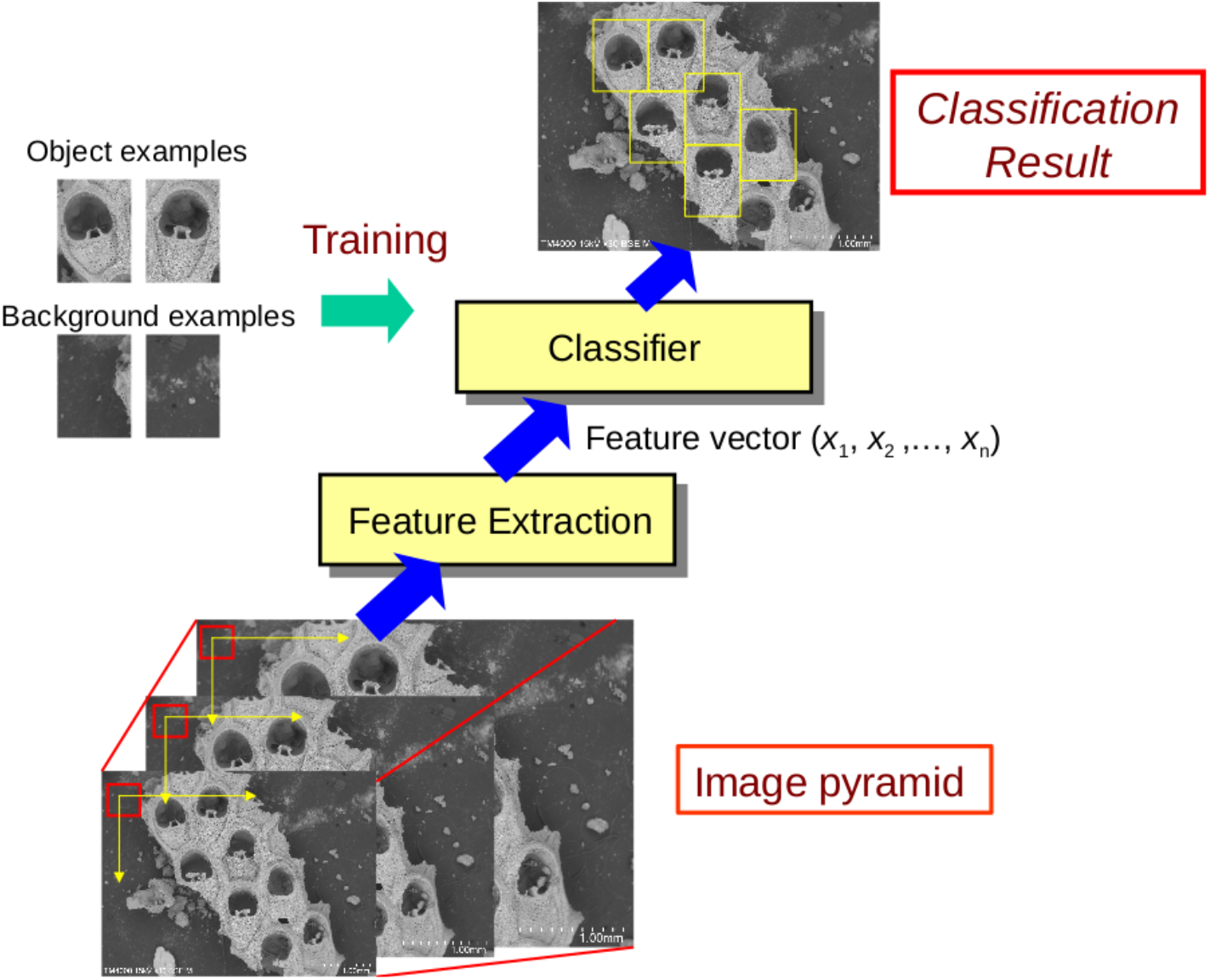
Diagram of object detection framework. In the proposed framework, a training set containing object and background examples is used to train a support vector machine (SVM) classifier. A sliding window is then scanned over an image pyramid and its features are then classified as either object or background. Detected objects are output by the model after non-maximum suppression of overlapping windows is performed.

### 2.5 Training shape predictors

We performed shape prediction by feeding objects detected using the above procedure to a cascade regression algorithm (Kazemi and Sullivan, 2014) that performs object-specific automated landmark detection. In brief, cascade shape regression predicts shape (S) in an iterative procedure using a sparse subset of pixel intensities collected on the area of the image where the object was identified. Starting with an initial shape ‘guess’ *S*_0_, the predicted shape (S) is refined through shape increments in a cascade of depth *K* (Figure 3), so that:

**Figure 3:**
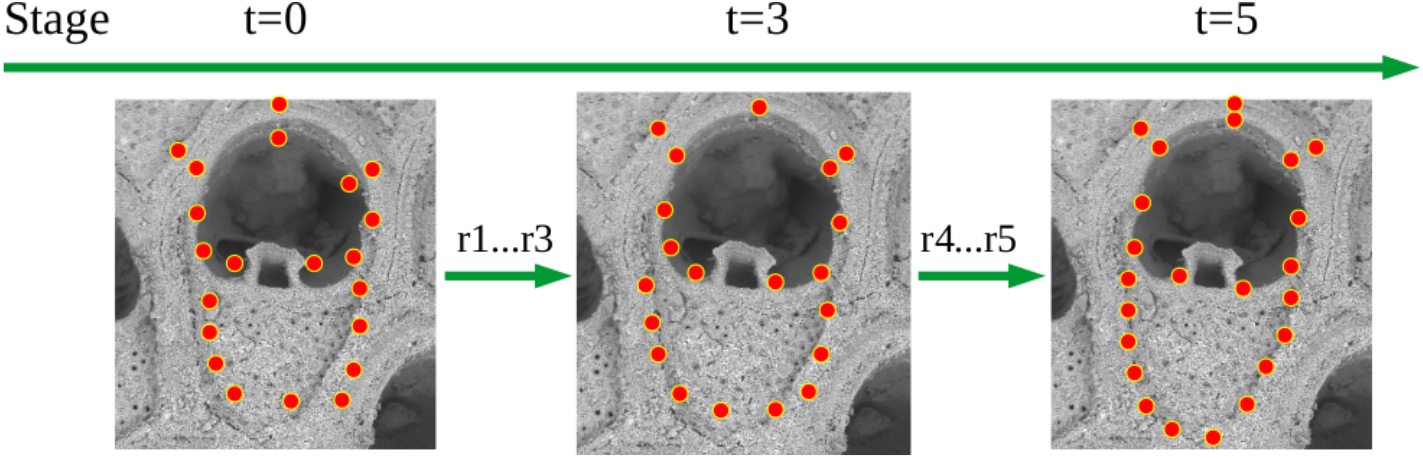
Diagram of shape prediction framework. In the proposed framework, a training set containing positional ground truths is used to train a cascade shape regression model. Shape prediction of unannotated objects is then performed using a sparse subset of pixel intensities collected on the area of the image where each object was identified. Cascade shape regression uses an iterative procedure (green arrows) to predict the shape of the object (orange landmarks). Starting with an initial shape ‘guess’ (landmarks at t=0), the predicted shape is refined through shape increments (using r regressors) in a cascade (from t=0 to t=5).

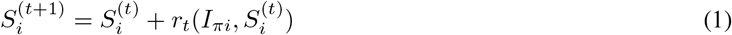

where *r*_*t*_ represents a trained regressor, *I_πi_* represents an image and 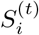 represents the previous stage’s shape estimate (Kazemi and Sullivan, 2014). Each regressor in the cascade is learnt using a gradient boosting algorithm with a square error loss function (Hastie, Tibshirani, and Friedman, 2009). The gradient boosting algorithm uses a user-defined number of regression trees of depth *T* for which decisions at each node are based on thresholding the difference in intensity values at a random pair of pixels, given an exponential prior over the distance between pixels used in a split. More details of the gradient boosting algorithm can be found in Kazemi and Sullivan (2014). During training, we performed data augmentation of the training set by adding in 300 random deformations of each training object, effectively boosting the number of training examples. Similar to object detection, we carry out an exhaustive grid search of the training parameters *T* (from 1 to 8) and *K* (from 10 to 30), and use the best performing model as the final model.

### 2.6 Testing object detectors – Precision and Recall

We evaluated the performance of object detectors using three metrics that are standard in the field: precision, recall and mean average precision (Powers, 2011). Precision is defined here as the ratio of true positive object predictions to all positive object predictions. Recall, also known as the true positive rate, refers to the ratio of true positive object predictions to all true positive objects present in the data. Finally, mean average precision is defined as the area under the recall-precision curve. All three performance measurements vary from 0 to 1 and can be directly compared across datasets. Note that the performance of the classifier on the test set will only correctly reflect the performance of the model on unlabeled data if these pictures were taken with the same general setup as in the training data (i.e., similar background and resolution), and not necessarily if image capture methods change dramatically (e.g., extremely low resolution images, such as 80×80).

### 2.7 Testing shape predictors – Euclidean distance

In order to quantify the accuracy in placement of landmarks, we measured the normalized Euclidean distance in pixels between each landmark’s location in the ground truth (test) set and as predicted by the model (Kazemi and Sullivan, 2014). The normalization process allows direct comparison of the error between the three datasets (and across studies), since image parameters are different in each study. Here, landmark distances were normalized by the total length of the structure. For the most complex dataset, we also break down the total measurement error by image file and landmark, allowing us to evaluate potential sources of error in landmark predictions.

### 2.8 Implementation and file conversion

All algorithms were implemented in Python using the following libraries: *numpy 1.13.3*, *pandas 0.22.0*, *dlib 19.7.0* and *opencv-python 3.4.0.12*. Our current implementation can be found at *https://github.com/agporto/ml-morph*. The scripts can be used out-of-the-box and can be run on any operating system. On the github page, we also provide a detailed vignette and an example dataset in order to illustrate the use of this software. Among the capabilities of this software, we should note that there is a preprocessing script that converts traditional landmark files from the most common morphometric packages (*tps* format; Rohlf, 2006) to standard input files used in training and testing of the object detectors and shape predictors used in our machine-learning phenotyping pipeline. This script allows previously landmarked datasets to be immediately used in automation.

## 3 Results

### 3.1 Object detection

The object detectors learnt based on HOG features achieved a high degree of recall and precision in all three datasets (Table 1). To a large extent, the training parameters had no major impact on the performance of the final model, with several different training parameter combinations resulting in similar performance (Figure 4). As expected, recall and precision were highest in the low and intermediate complexity datasets (100%), and slightly lower in the high complexity dataset (95%). Using the high complexity dataset as a benchmark, we can infer that false positives occur at low frequency (5%; Table 1) and can be effectively eliminated by a simple size filter without incurring in loss of true positives. For example, eliminating zooids with size smaller than 40% of the mean population size in the highly complex dataset leads to the elimination of all false positives, without eliminating true positives.

**Table 1:** Training and testing parameters of the best fit object detection model. In this table, we detail the training parameters of the best fit object detection model for each of the landmark datasets used in this manuscript, together with their performance on the test set. Precision is defined here as the ratio of true positives to all positives. Recall refers to the ratio of true positives to all true positives. Mean average precision is defined as the area under the recall-precision curve. Details of the training parameters can be found in the main text and in (Powers 2011)..

| Dataset | Train |  |  | Test |  |  |  |
| --- | --- | --- | --- | --- | --- | --- | --- |
| | N | C | $\epsilon$ | N | Precision | Recall | MAP |
| Low | 224 | 1 | $10^{-4}$ | 56 | 100% | 100% | 100% |
| Intermediate | 144 | 3 | $10^{-2}$ | 36 | 100% | 100% | 100% |
| High | 320 | 5 | $10^{-2}$ | 80 | 95.4% | 95.4% | 95.1% |

N = sample size; C = soft margin parameter ; $\epsilon$ = penalty parameter ; MAP = mean average precision

**Figure 4:**
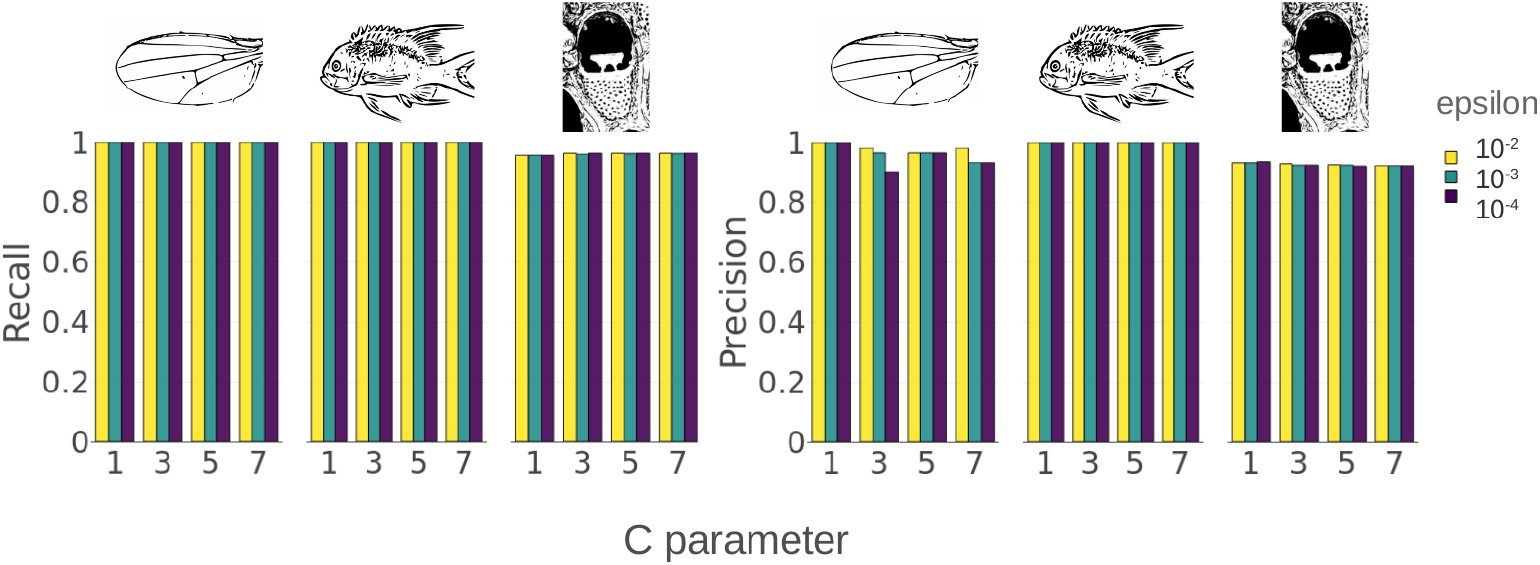
Grid search for optimal object detection training parameters. Barplots illustrating the recall and precision obtained by object detection models trained with varying training parameters. Each dataset is represented by a drawing of an example specimen. In our case, we varied the soft margin parameter C (X-axis, from 1 to 7) and the insensitivity zone (colored bars, from 10^*−*2^ to 10^*−*4^). The C parameter regulates the distance between the hyperplane that separates the two classes and the closest data point. The insensitivity zone (*ϵ*) regulates the penalty associated with errors in classification.

### 3.2 Shape prediction

We report shape prediction results for the three datasets in Table 2. The table contains comparisons of the magnitude of the observed error of the best fit model against the theoretical minimum (intra-observer error) and against the best performing semi-automatic methods available in the literature. All errors are reported as normalized mean Euclidean distances. The magnitude of error is low across all datasets. In the low complexity dataset, the best-fit model presents a normalized error of 0.51% of the wing length (2.8 pixels in raw values). This magnitude of measurement error is remarkably close to the theoretical maximum accuracy that is possible given intra-observer measurement error (0.34% or 1.8 pixels). The best-fit model performs equivalently to the best semi-automatic approach available in the literature (0.67% or 3.54 pixels; Loh et al., 2017 -Table 5) and considerably better than others (e.g., >2% or > 10.6 pixels for Houle et al., 2003, as reported in Loh et al., 2017-Table 5). Search of the training parameter space reveals that similar results can be obtained for different tree depths and cascade depth (Figure 5). In other words, prediction performance is generally high, regardless of model fine-tuning. Relatively small models (low depths) still provide robust results. Over-fitting can only be observed for tree depths larger than six.

**Table 2:** Training and testing parameters of the best fit shape prediction model. In this table, we detail the training parameters of the best fit shape prediction model for each of the landmark datasets used in this manuscript, together with their performance on the test set. Error is measured here as the normalized Euclidean distance. The literature tab refers to the performance of the best semi or fully-automatic model present in the literature for each study system. Given that drosophilids are the only structure that was automated in the past, only results for drosophila algorithm are detailed. Details of the training parameters can be found in the main text and in Dalal and Triggs (2005) and King (2009).

| Dataset | Train |  |  |  | Trees (N) | Feature pool size | Test splits | Oversampling(N) |
| --- | --- | --- | --- | --- | --- | --- | --- | --- |
|  | N | nu | T | K |  |  |  |  |
| Low | 224 | 0.1 | 3 | 30 | 500 | 1200 | 20 | 300 |
| Intermediate | 144 | 0.1 | 2 | 30 | 500 | 1200 | 20 | 300 |
| High | 320 | 0.1 | 2 | 25 | 500 | 1200 | 20 | 300 |

| Dataset | Test |  |  |  |
| --- | --- | --- | --- | --- |
|  | N | Error | Minimum | Literature |
| Low | 56 | 0.513% | 0.330% | 0.6%(Loh et al. 2017); >2% (Houle et al ,2003)* |
| Intermediate | 36 | 0.810% | 0.540% | None |
| High | 80 | 2.090% | 1.310% | None |

N = sample size; nu =regularization parameter; T= tree depth; K = cascade depth
'\*' as reported in Loh et al. (2017) .

**Figure 5:**
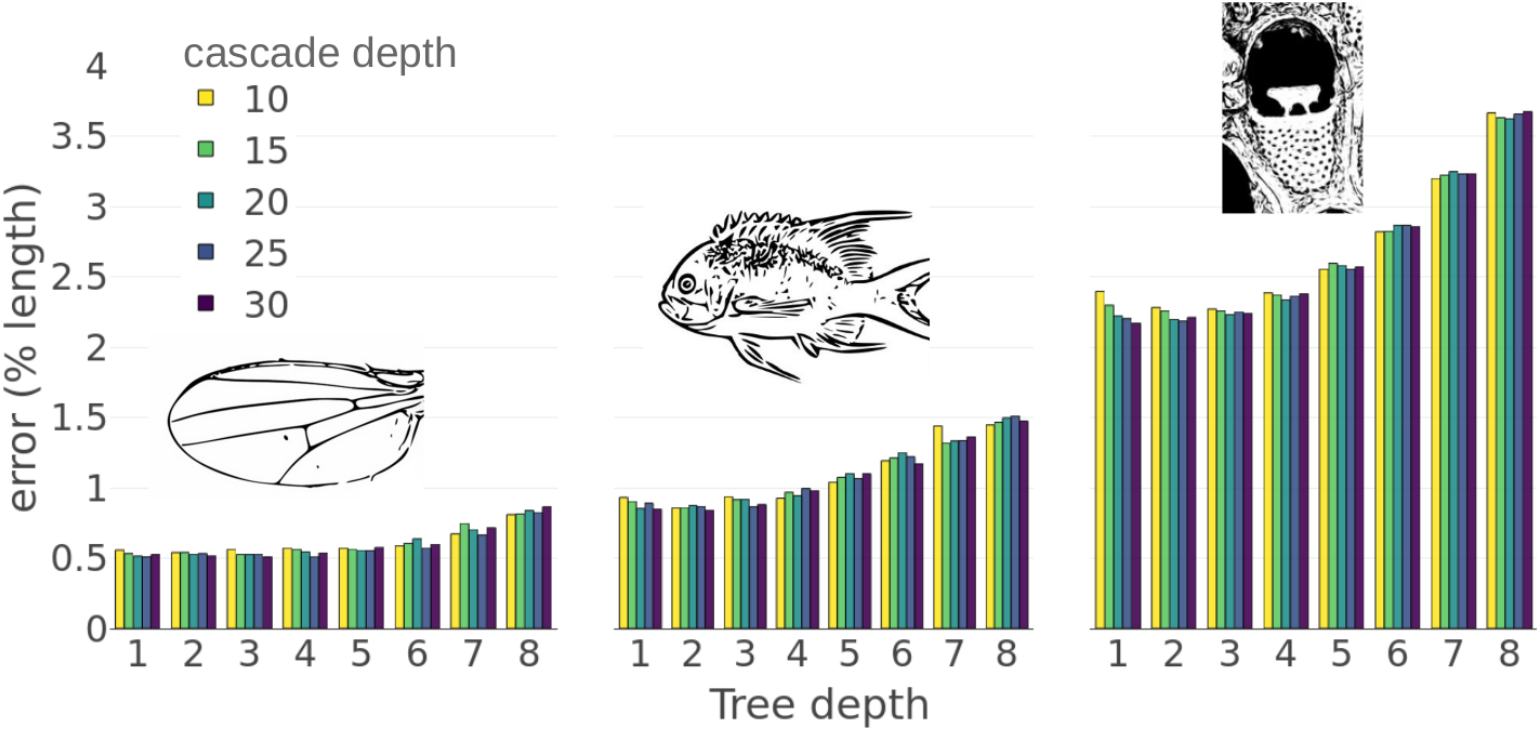
Grid search for optimal shape prediction training parameters. The barplot illustrates the normalized mean error obtained by shape prediction models trained with varying training parameters. Each dataset is represented by a drawing of an example specimen. In our case, we varied the regression tree depth (X-axis, from 1 to 8) and the cascade depth (colored bars, from 10 to 30). Tree depth refers to the depth of the decision trees used by the gradient boosting algorithm. The cascade depth refers to the number of shape updates that are used to predict each object’s shape, given local pixel patterns.

While the degree of error observed for the intermediate complexity dataset is larger compared to the low complexity dataset (0.81% or 4.8 pixels, Table 2), it is still similar to the intra-observer error (0.54% or 3.2 pixels). Search of the training parameter space reveals that greater tree depths lead to over-fitting and pronounced performance loss (Figure 5). The depth of the cascade has only a mild impact on the results.

Finally, the degree of error observed for the high complexity dataset is higher than for the other two datasets (2.1% or 6.2 pixels, Table 2), in a large part because intra-observer error is also higher (1.3% or 3.8 pixels). Notably, the ratio of the best-fit model error relative to intra-observer error is remarkably similar across all datasets, with the shape predictors producing datasets with a degree of error 50% higher than the theoretical minimum. While 50% might seem like a substantial difference, note that in drosophilids this is equivalent to a difference of one pixel.

Closer examination of the sources of error in the complex dataset reveals two important sources of error in our approach. First, mean average error varies from 1.1% to 3.5% across image files (Figure 6). Differences among image files are associated with differences in the quality of preservation of fossil specimens and with the quality of image capture. Figure 6 illustrates one of the best and worst performing images, which vary considerably in the level of contrast and taphonomy. Similarly, accuracy in landmark predictions varies significantly across landmarks, with certain landmarks presenting mean average error around 0.91%, while others present values of 4.2% (Figure 7). The presence of clear edges and high contrast is clearly associated with accuracy in the predictions (Figure 7).

**Figure 6:**
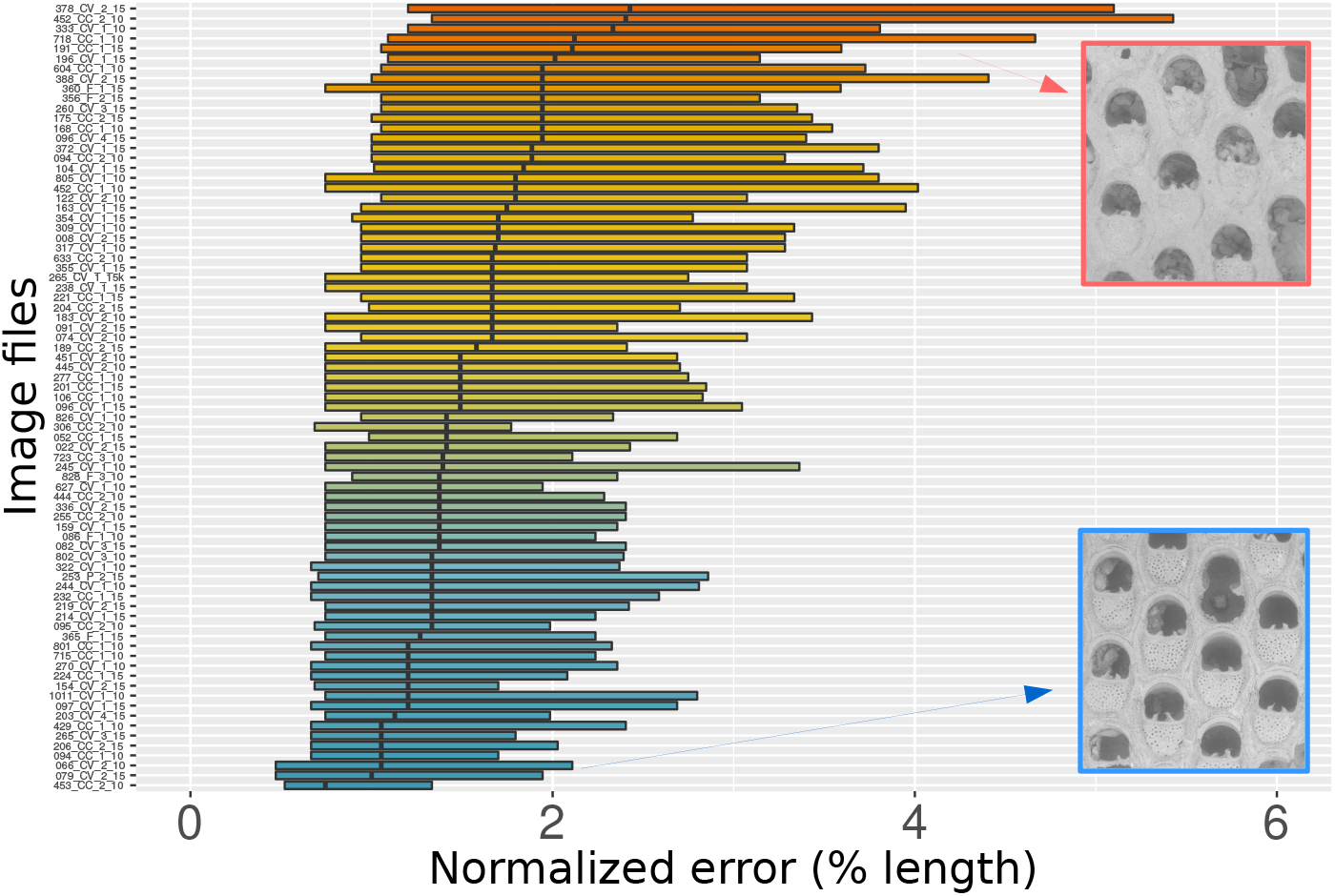
Breaking down the prediction error according to image quality. Each boxplot illustrates the distribution of landmark prediction errors of our best fit model on each image of the test set (whiskers omitted for clarity). Image files are sorted according to their median prediction error, from low (blue) to high (orange). Representative specimens from the extremes of the distribution are shown near the upper and lower margins. Note the difference in fossil degradation and image contrast between the two specimens. Note also that images with poorer image quality (orange bars) have more variance in prediction error among detected objects.

**Figure 7:**
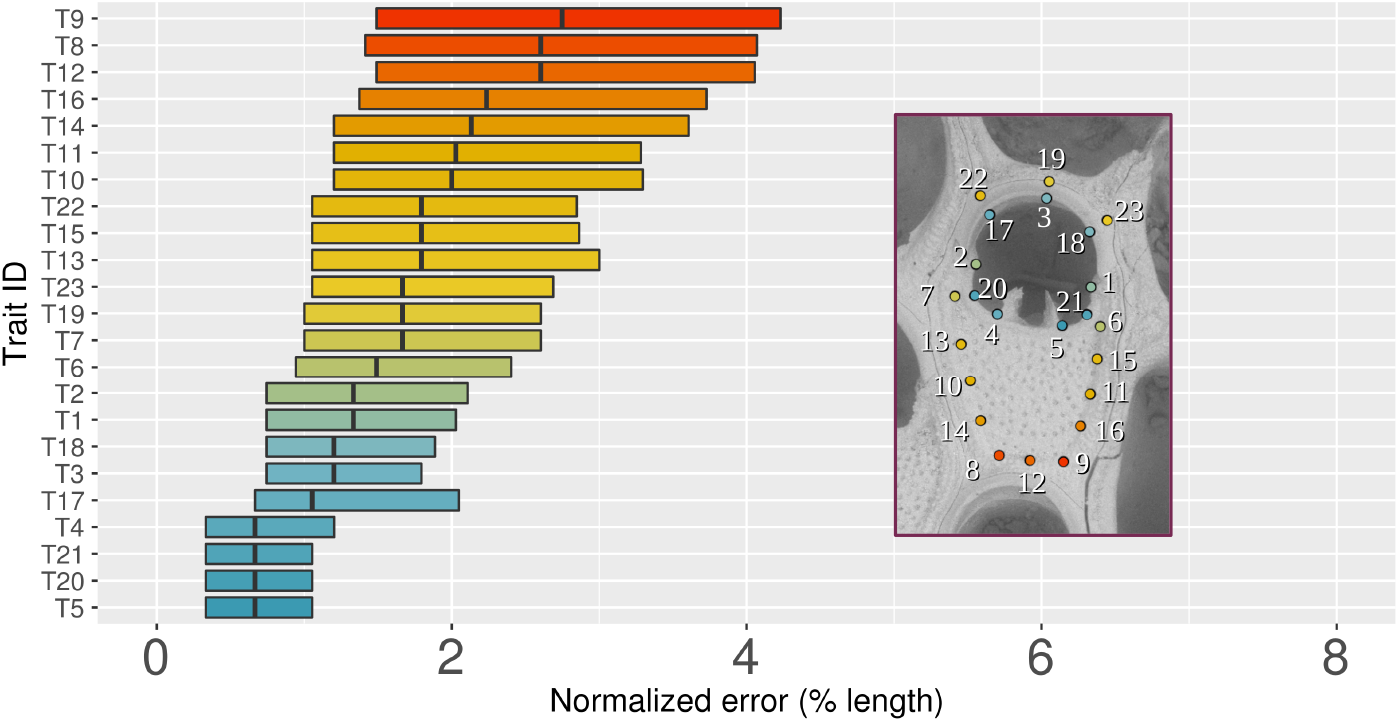
Breaking down the prediction error among the landmarks. Each boxplot illustrates the distribution of landmark prediction errors of our best fit model according to each landmark (whiskers omitted for clarity). Landmarks are sorted according to their median prediction error, from low (blue) to high (orange). We also show one representative specimen with landmarks colored according to the degree of prediction error. Note the difference in the degree of error between landmarks in low (e.g. T8, T9) and high (e.g., T4, T5) contrast areas. Note also that the variance in the distribution of errors per landmark is correlated with the median.

### 3.3 Implementation speed

We used the high complexity dataset to benchmark the speed of the pipeline, when run on an Intel Core i7 2.7Ghz with 16GB of RAM. Using the training parameters of the best fit models (Tables 1 and 2), object detector training occurred in 9.3 minutes per run, and testing took a total of 15 seconds per run. Shape predictor training occurred in 9.23 hours per run, while testing took only 7 seconds per run. To a large extent, the amount of oversampling applied to the training set is the key parameter regulating the training times for shape predictors, with higher oversampling amount requiring longer training.

In order to benchmark the speed at which the pipeline can generate predictions for new images, we applied the final object detector and shape predictor to a larger sample of *Steginoporella* images (N=1684). In this larger sample, the entire automated landmarking pipeline took 3.12 minutes to process all images, during which it identified 23 landmarks per zooid in a total of 13,686 zooids at a resolution of 800 x 600 pixels. Based on generous assumptions (3 minutes per zooid), an experienced morphometrician would spend 684 hours (about 85 eight-hour uninterrupted workdays) to collect a comparable dataset.

## 4 Discussion

Morphometric characterization of biological structures has become an essential component in studies of morphology. However, morphometric data collection has remained mostly manual, in large part due to the lack of a general framework for automation (see Hsiang et al., 2018 for a notable exception). In this study, we propose a simple, fast and accurate pipeline for automation of landmark collection in any semi-rigid biological structure. Our approach is based on supervised learning and uses a combination of object detection (Dalal and Triggs, 2005) and shape prediction (Kazemi and Sullivan, 2014) to accurately place landmarks of interest on one or several objects in an image.

### 4.1 Model performance

Object detectors performed well in all three datasets, making no mistakes in the low and intermediate complexity datasets (Table 1, Figure 4). Even in the most complex dataset consisting of pictures of fossil specimens in various taphonomic states, object detection rates were high (95%), indicating detection should not be a concern for the standardized images datasets typically analyzed in biological studies. Note, however, that the high performance of object detectors in the low and intermediate complexity datasets is due to the high standardization of image capture method and should not be used as the null expectation for less standardized data.

As expected, the shape prediction part of the pipeline is where differences in accuracy among data sets were the largest (Table 2, Figure 5). In drosophilid wings, accuracy is homogeneous across landmarks and very similar to the scale typically found among human observers. In *Steginoporella*, heterogeneous levels of accuracy depend on the landmark and the individual specimen (Figure 6 and 7). We highlight two main sources for the heterogeneity in accuracy in the *Steginoporella* dataset. First, image quality varied considerably across specimens, largely due to fossil degradation. Since the *Steginoporella* samples are composed exclusively of fossil specimens (see Liow et al., 2017), some of which were collected at harsher depositional environments than others, some specimens present a higher degree of damage and cementation than the remaining ones, leading to a reduction in contrast between the morphological features of interest and the background. Additionally, some landmarks are placed in contrast poor locations (Figure 6), adding noise to automation based on HOG features, as this is informed by local image contrast (Dalal and Triggs, 2005).

### 4.2 Advantages of the framework

Given the relative small size of our training sets and how well the pipeline performs on datasets of different complexity, we argue here that this pipeline has general applicability in biology. The main advantages of this framework are its accuracy, speed, and infrastructure requirement. Even in the most complex dataset, the landmarking pipeline being proposed here exceeds the accuracy reported in many system-specific landmark approaches observed in the literature (Loh et al., 2017; Houle et al., 2003). Note also that the intra-observer error, that we define as the theoretical minimum error when evaluating our method, would probably be larger if multiple people had annotated such datasets, as commonly done in morphometric studies. Furthermore, potential errors committed by the pipeline can be manually corrected after prediction using the image software of choice (e.g., imglab (King, 2009) and/or tpsDig (Rohlf, 2006)). When comparing time-weighed performance with manual annotation, the benefits of automation are even clearer.

Manual annotation of the entire *Steginoporella* dataset would take around 85 days, using generous assumptions of the work-load (3 minutes per zooid). In about the same amount of time (3.12 minutes), our pipeline is able to process 1,684 images, representing 13,686 zooids and 23 landmarks per zooid. This time can even be reduced to 1.5 minutes if a lower image resolution is used, though that is likely to increase error levels. Moreover, if a previously annotated dataset has already been collected, the whole procedure, from producing the training sets to training the object detection and shape predictors to obtaining an automated landmarking framework, can be performed on a personal computer in the timespan of two workdays. The proposed methodology for object detection, while not in our purview, can also be used for other purposes in biology. For example, based on the detected objects, one could adapt this framework to count objects and to obtain, for example, the spacing of objects on an image, such as the distance between the zooids of *Steginoporella*.

In our view, this automated landmarking framework opens up the possibility of development of truly high throughput high-dimensional phenotyping procedures, which will propel biology into the age of phenomics (Houle, Govindaraju, and Omholt, 2010).

### 4.3 Limitations of the framework

The most important limitation in the proposed pipeline is that HOG based object detectors are partially sensitive to changes in orientation (Dalal and Triggs, 2005). As a consequence, objects cannot be detected when upside down, for example. In our view, these limitations can be overcome using one of the following: (1) training of multiple object detectors, one for each position or (2) through the use of CNN-based detectors, such as Ren et al. (2015). While CNN based detectors are insensitive to changes in orientation, it is worth pointing out that such algorithms require a larger training dataset (more images), higher hardware specifications, are less portable (file size), and require more fine tuning (Suleiman et al., 2017).

Another limitation of the proposed pipeline is that it can only extract data from 2D images. A non-trivial portion of geometric morphometrics is done in 3D, and while the techniques presented here could be expanded to 3D objects, we currently do not have an efficient implementation of it. At most, our proposed framework can be applied to 2D slices of 3D structures, but while this might prove useful in some systems (Hsiang et al., 2018), it cannot be considered an explicit 3D approach. Finally, although we provide python code that can be used out-of-the-box, we strongly recommend that other authors explore all training parameters for object detection and shape prediction, as those can have significant impact on the accuracy of the final model (Kazemi and Sullivan, 2014). This is especially true if image parameters (e.g., resolution) are significantly different from the three datasets presented here.

## 5 Conclusions

We have developed a machine-learning pipeline (ML-morph) for automated detection and landmarking of biological structures in images, which can be used to collect morphometric data at a large scale. ML-morph opens up the number of possibilities for automation within the morphometric community, greatly increasing the scale of the questions being asked and opening up new research avenues that previously faced sample size barriers.

## Supporting information

Supporting Information 1 - Drosophilid IDs from Morphbank

## 6 Acknowledgements

AP and KLV were funded by the The Research Council of Norway — grant no. 249961. The authors wish to thank Mali Ramsfjell for helping with the *Steginoporella* sample preparation and imaging, and Lee Hsiang Liow for access to the scanning electron microscope (TM-4000). The scanning electron microscope (TM-4000) used in this study has been funded by The European Research Council — grant no. 724324.

## 7 Author’s contribution

AP and KLV conceived the project; AP collected the data and designed the methodology; AP analyzed the data and wrote the code; All authors contributed to writing the manuscript, with AP having the lead on the writing.

